# Structural insights into interaction mechanisms of alternative piperazine-urea YEATS domain binders in MLLT1

**DOI:** 10.1101/836932

**Authors:** Xiaomin Ni, David Heidenreich, Thomas Christott, James Bennett, Moses Moustakim, Paul E. Brennan, Oleg Fedorov, Stefan Knapp, Apirat Chaikuad

**Affiliations:** Institute of Pharmaceutical Chemistry, Goethe-University Frankfurt, 60438 Frankfurt, Germany; Structural Genomics Consortium, BMLS, Goethe-University Frankfurt, 60438 Frankfurt, Germany; Target Discovery Institute and Structural Genomics Consortium, University of Oxford, Oxford OX3 7DQ, UK

**Author notes:** C4 Therapeutics Inc., 490 Arsenal Way, Suite 200, Watertown, MA 02472.

## Abstract

YEATS-domain-containing MLLT1 is an acetyl/acyl-lysine reader domain, which is structurally distinct from well-studied bromodomains and has been strongly associated in development of cancer. Here, we characterized piperazine-urea derivatives as an acetyl/acyl-lysine mimetic moiety for MLLT1. Crystal structures revealed distinct interaction mechanisms of this chemotype compared to the recently described benzimidazole-amide based inhibitors, exploiting different binding pockets within the protein. Thus, the piperazine-urea scaffold offers an alternative strategy for targeting the YEATS domain family.

---

Epigenetic signaling plays crucial functions in chromatin-dependent transcription, and often requires tight regulation involving post-translational modifications essentially at specific lysines of several key proteins, including histone proteins. Recognition of acetylated lysine, one of the most common epigenetic marks, is one of the hallmark events in epigenetic signaling. In addition to the well-established readers such as bromodomains and some double PHD finger domains^1–2^, YEATS domain, which have been described in four human proteins including MLLT1 (ENL), YEATS2, MLLT3 (AF9) and glioma amplified sequence 41 (GAS41 or YEATS4), has been classified as a protein class capable of recognizing histone acetylation as well as bulkier lysine posttranslational modifications such as propionylation, butyrylation, crotonylation and succinylation^3–9^. YEATS-domain-containing proteins have potentially distinct preferences towards different lysine modifications on histone proteins. For instance, MLLT1/3 prefer acetylated Lys9, Lys18 and Lys27 of histone H3^4, 10^, while YEATS2 has been demonstrated to have a stronger affinity for crotonylation marks^11–12^. In addition to moderate affinities for acetyl-lysine, some unusual reader activities of GAS41 have been reported including the recognition of histone succinylation in a pH-dependent manner^13^, as well as its ability to form a dimer through the C-terminal coiled-coil domain for cooperatively bivalent binding to di-acetylated histone^14^.

Several reports have associated YEATS-domain-containing proteins as a key driver for the development of diseases, in particular cancer^3^. For examples, chromosomal rearrangements involving the MLLT1 as well as MLLT3 YEATs domains promote development of leukemia ^10, 15–16^. The YEATS2 gene is highly amplified in non-small cell lung cancer (NSCLC), and its role regulating transcription is essential for tumorigenesis^17^. In addition, overexpression of growth-promoting GAS41 has been associated with malignancy of many cancer types, including gastric^18^ and hepatic carcinomas^19^ as well as NSCLC^20^. These consistent lines of evidence have suggested therefore YEATS domains as a potential target for chemotherapeutic treatment.

YEATS domains exhibit highly distinct beta-sheet topology compared to other acetylation readers such as helical bromodomains. The YEATS acetyl-lysine binding site comprises a more open, shallow, surface-exposed binding channel, which is open on both sites allowing accommodation of larger lysine modifications^4, 21^. These distinct structural features suggest that YEATS domain inhibitors require different structural features compared to acetyl-lysine mimetic groups explored for the development of bromodomain inhibitors^22–23^. An early attempt exploiting a peptide-based strategy with 2-furancarbonyl lysine led to a competitive nanomolar binder^24^, demonstrating potential inhibitions of the YEATS protein module. Our recent screening efforts alternatively revealed a number of small molecule binders for MLLT1/3^21, 25^. One chemotype identified was ben-zimidazole-amide, and extensive modification focused on this scaffold led to the development of the potent and selective MLLT1/3 inhibitor (SGC-iMLLT) with IC_50_ of ~0.26 μM and *K*_D_ of 113 nM^26^. Unfortunately, moderate metabolic stability of this SGC-iMLLT chemical probe has limited its further use in chemotherapeutics. Nevertheless, the discovery of this chemical probe has demonstrated therefore a potential of targeting YEATS domains using small molecule inhibitor.

Alternative MLLT1/3 binders with different chemical moieties may offer a benefit in cellular activities of MLLT1 inhibitors. We therefore aimed to search for other scaffolds distinct from benzimidazole-amide, which is the only characterized binder to date. Analyses of the results from our previous screening campaign revealed a number of piperazine-1-carboxamide hits^25^, suggesting that this piperazine-urea scaffold could serve an alternative acetyl-lysine mimetic moiety for MLLT1/3. Interestingly, this chemical class has been extensively used previously in development of inhibitors with good pharmacokinetic properties for diverse targets, such as fatty acid amide hydrolase (FAAH)^27^, melanocortin subtype-4 receptor (MC4R)^28^ and the receptor for the chemokine CCL2 (CCR2 or known also as MCP-1)^29^. For the potential application as an alternative scaffold for YEATS domain inhibitors, we investigated interaction mechanisms of the piperazine-urea chemotype in MLLT1.

Based on our previous work^25, 30^, we selected three piperazine-1-carboxamide hits (**1, 2** and **3**) and for comparison three benzimidazole-amide derivatives of similar sizes (**4, 5** and **6**), and first determined the crystal structures of their complexes with MLLT1 (Figure 1). We observed surprisingly different binding modes and interactions between these two chemotypes to the protein. The most remarkable change was the orientation of their central primary amide cores. This functional group of the piperazine-urea derivatives flipped horizontally in comparison to that of the benzimidazole-amide compounds, and instead assumed a similar binding orientation to that of acetyl-lysine^21^. Nevertheless, similar β-sheet type hydrogen bonding patterns between the amide carbonyl atom to the backbone amine of Tyr78 as well as contacts with the conserved water molecule and Ser58 were highly maintained. Comparable binding modes between the secondary amide central core of the piperazine-1-carboxamide compounds and acetyl-lysine confirmed our previous hypothesis on the use of the amide group as an acetyl-lysine mimetic moiety for YEATS domains^21^.

**Figure 1.**
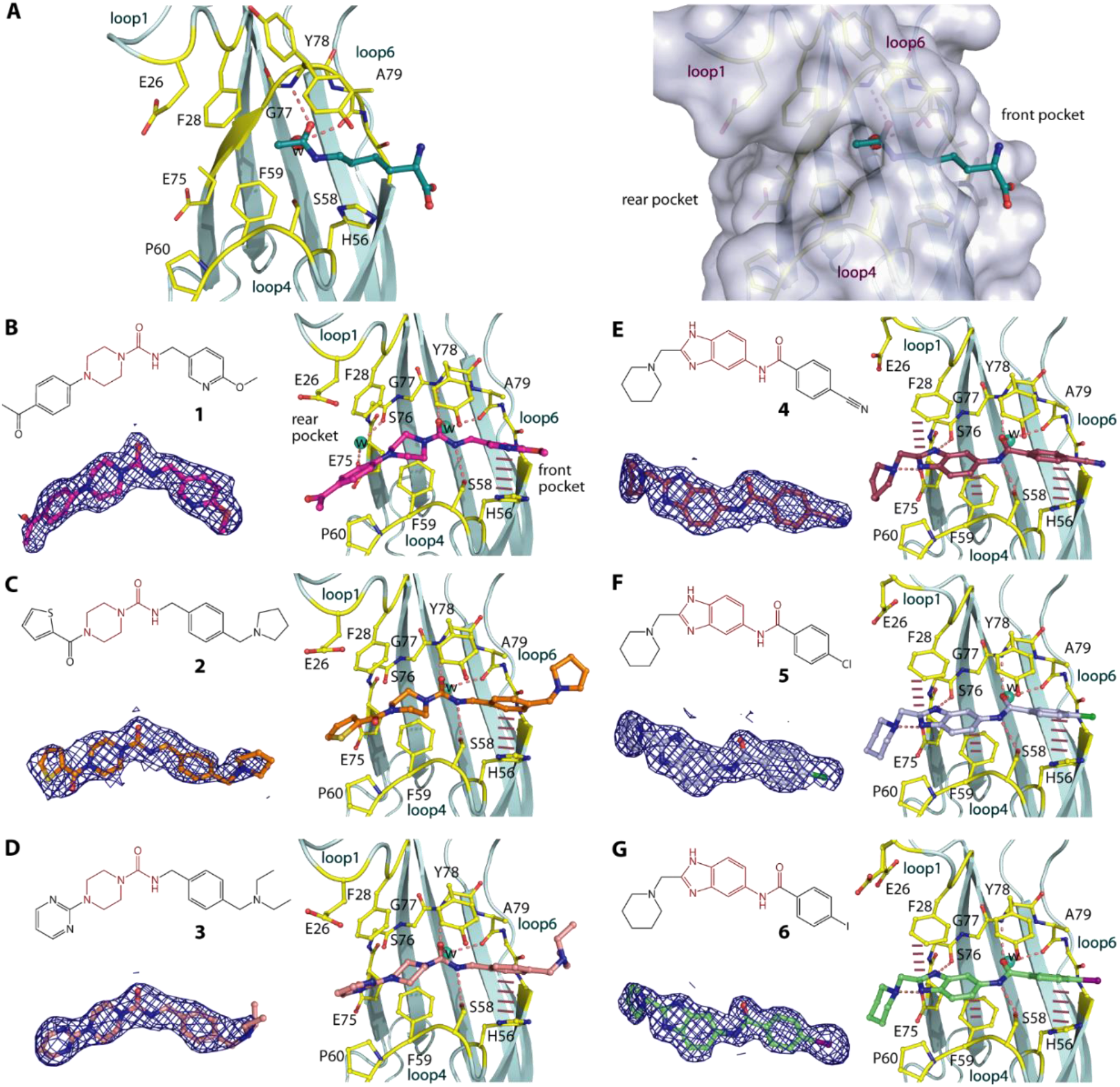
Piperazine-urea and benzimidazole-amide derivatives and their binding modes in MLLT1. **A**) The binding mode of acetyllysine in MLLT1 (pdb id: 6hpz) revealing potential front and rear pockets that could be targeted by small molecules. Chemical structures and the binding modes in MLLT1-complexed crystal structures of three piperazine-1-carboxamide **1, 2** and **3** (**B-D**; pdb ids: 6t1i, 6t1j and 6t1l) and three benzimidazole-amide **4, 5** and **6** (**E-G**; pdb ids: 6t1m, 6t1n and 6t1o). Dashed lines indicate hydrogen bonds, while bar, dashed lines are for potential aromatic π-stacking. Water molecules are shown in spheres. Electron density maps contoured at 1σ for the bound ligands are shown in the insets below the chemical structures in each panel. See supplementary figure S1 for omitted electron density maps.

Earlier crystal structures suggested an important role of aromatic moieties on both ends of the amide central core for achieving aromatic stacking with Phe28, Phe59 and His56^21, 30^, as exemplified also here in the interactions of **4, 5** and **6** (Figure 1E-G). However, in contrast to benzimidazoles, all piperazine-urea **1, 2** and **3** did not follow this scheme, and contained instead an sp^3^ piperazine on the *C*-linked region of the amide, whereas a benzyl substituent featured on the *N*-linked portion of the amide. This resulted in slightly different accommodation of their decorations within the protein (Figure 1B-D). At the front pocket, despite elongated by one carbon spacing the phenyl moiety of the benzyl ring was still located in proximity to His56 for aromatic stacking with this residue. Apart from this, there was no direct contact with the protein observed for the decorations at the other rear end. Although sharing the same binding pocket as the phenyl part of benzimidazole, the lack of aromaticity of the piperazine moiety restricted a *π*-stacking contact with Phe59. In addition, the orientation of the amide core in a canonical acetyl-lysine binding mode forced the aromatic decorations on the piperazine ring at the 4-position to assume a different trajectory than that of observed for the benzimidazole counterparts, and to occupy otherwise space adjacent to loop 4, which was not exploited previously by others MLLT1 binders. However, this binding mode diminished a *π*-stacking contact with Phe28 and feasibly resulted in more solvent-exposed for these extended groups.

Structural comparison between the piperazine-1-carboxamide- and benzimidazole-amide-MLLT1 complexes revealed that in addition to different binding modes of the compounds there were a number of conformational rearrangements within the protein (Figure 2A-B). Consistent with our previous observation for fragment binding^21^, the flipped amide with different binding positions likely affected slight adjustment of Ser58 side chain, which was required for maintaining their contacts. More dramatic structural alteration was evident for loop 1, which was feasibly a consequence of different ligand binding at the rear pocket. In comparison, the ‘closed’ conformation of loop 1 in the piperazine-1-caboxamide-MLLT1 complexes remained similar to that in the apo form^21^, albeit with a slight shift of Phe28 side chain likely due to steric effect from the piperazine moiety (Figure 2A). This suggested that the space adjacent to loop 4 might typically exist in the ligand-free state, and binding of the decorated piperazine moieties within this cavity did not significantly perturb structural integrity of the protein. In contrast, the presence of the benzimidazole ring within a cavity located in the rear pocket towards β5, which was filled by water in the MLLT1-**1** complex, led to significant structural alterations. This was evident not only for ~34° twist of Phe28 side chain for an optimal aromatic stacking, but an outward movement of loop1 by ~3-4 Å, adopting an open conformation (Figure 2B). These structural alterations re-modelled the shape of the rear pocket, which nevertheless could accommodate both ligands (Figure 2C). The binding modes of the piperazine-urea and benzimidazole-amide derivatives revealed therefore not only cavities that can be targeted for the design of more potent ligands interacting with the rear pocket, but they demonstrated also significant plasticity of the MLLT1 pocket.

**Figure 2.**
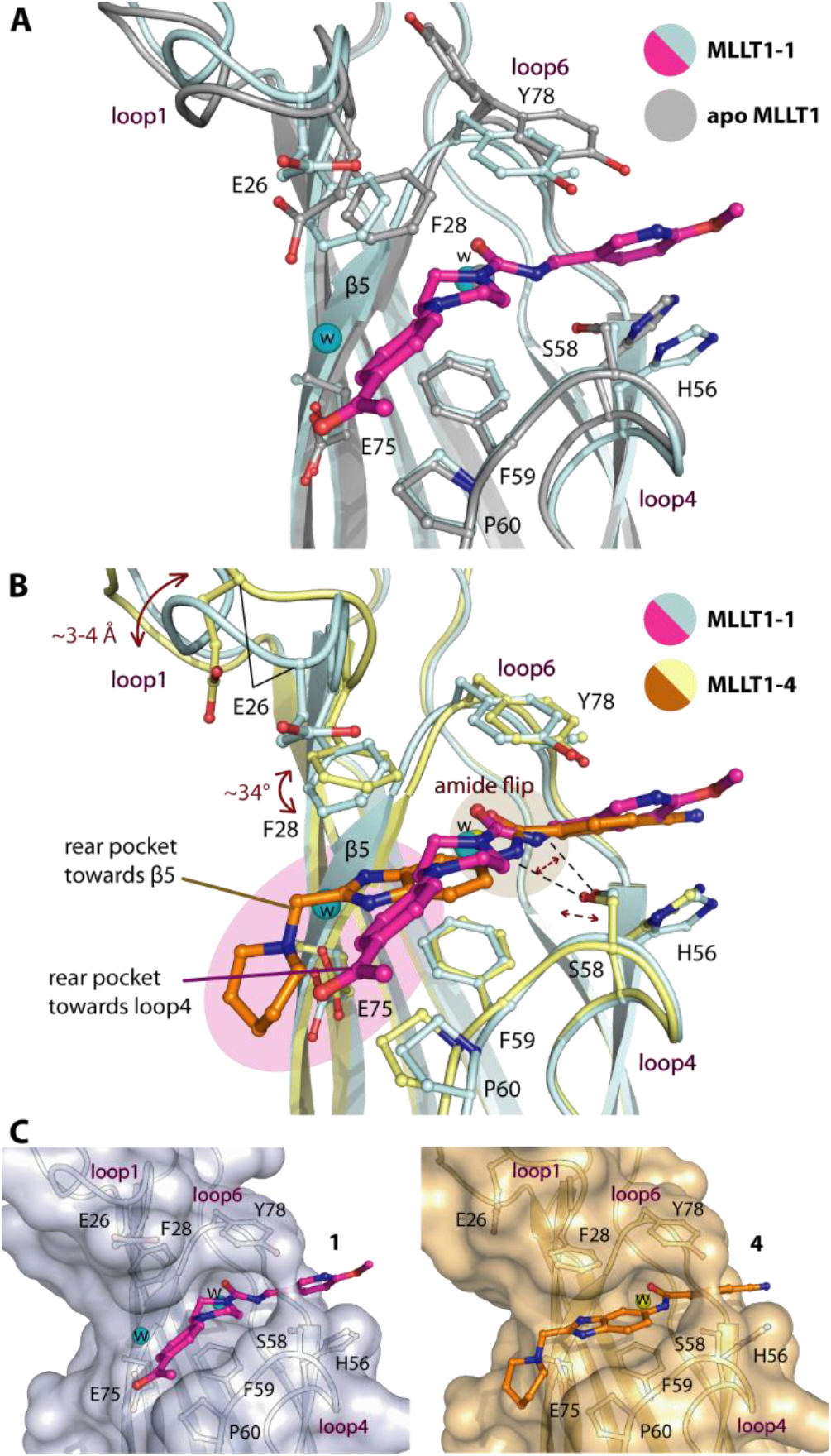
Structural comparison of interaction mechanisms of piperazine-1-carboxamide **1** and benzimidazole-amide **4** in MLLT1. A) Superimposition of the apo (pdb id 6hq0) and the **1-**complexed structures demonstrates no significant conformational changes for MLLT1. B) Comparison of **1-** and **4**-complexed structures reveals alterations of loop1, Phe28 as well as Ser58 potentially due to the orientation of the amide central core and different binding modes of the ligands that occupies different space at the rear pocket. C) Surface representation shows that conformational changes of MLLT1 adjust the shapes of the binding pocket complementarily for different compounds.

We next assessed binding affinities of all six derivatives for MLLT1, and performed isothermal titration calorimetry (ITC) to analyze binding constants and thermodynamics. All compounds exhibited binding potencies in a micromolar range (Figure 3A and B). Among the piperazine-urea derivatives, **1** was the most potent binder with *K_D_* of ~5.5 μM, which was ~3-fold lower than those of **2** and **3**, and was comparable to those of benzimidazole counterparts that showed also low micromolar affinities. Interestingly, we observed also that the trend of unfavorable entropy was characterized for the binding of all piperazine-urea compounds, which was in contrast to a gain in entropic contribution measured for benzimidazole-amide inhibitors. This change in thermodynamic signatures was in agreement with their diverse interaction mechanisms. However, the thermodynamics parameters observed from moderate binding in ITC remained highly estimated, and therefore further experiment might be required to confirm such differences.

**Figure 3.**
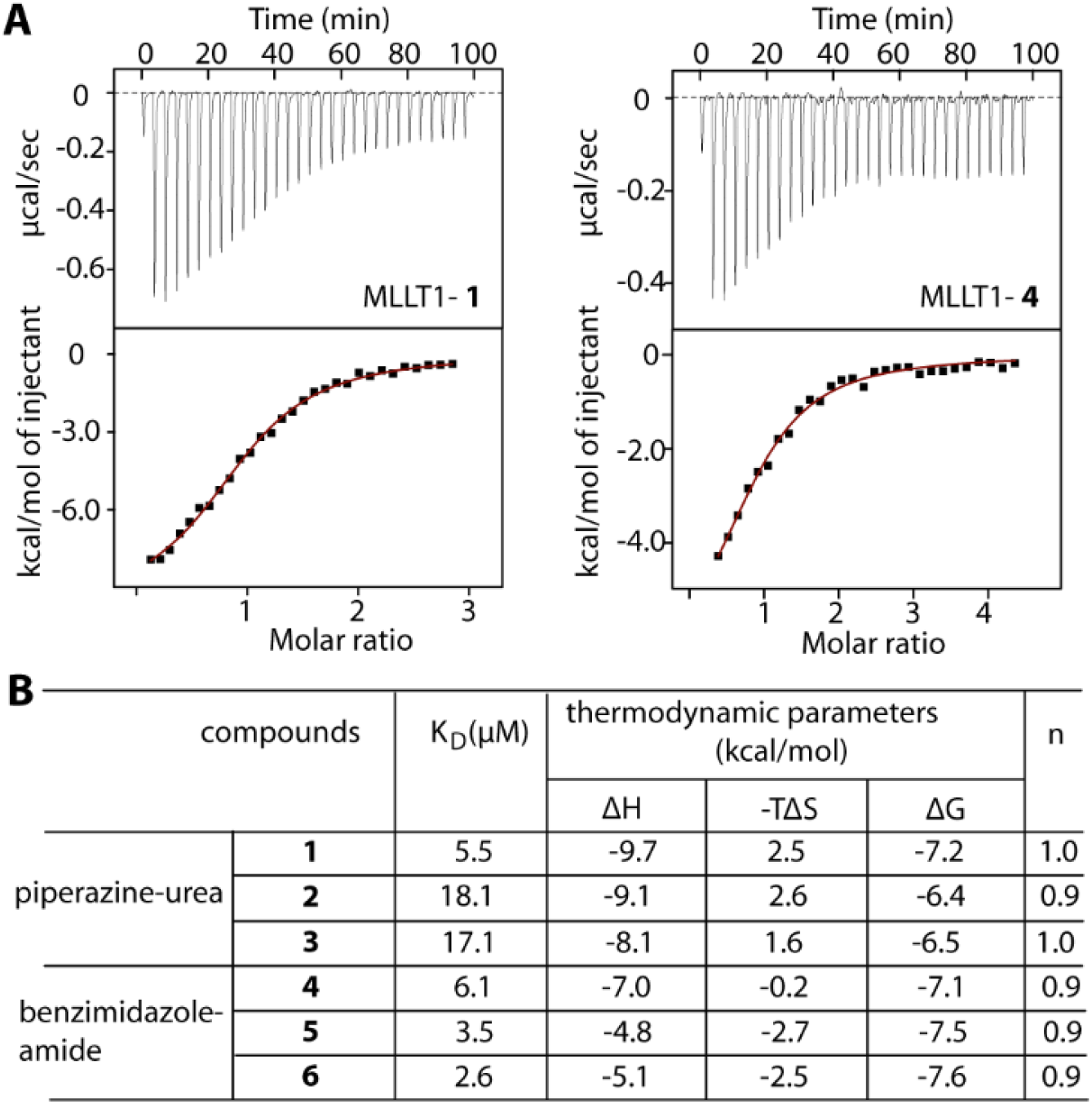
Binding of the derivatives in MLLT1 measured by ITC. A) Examples of ITC data for **1** and **4**. Shown are the isotherms of raw titration heat (top) and the normalized binding heat with the single-site fitting (red line; bottom). B) Summary of binding constants and thermodynamic parameters.

We next performed orthogonal AlphaScreen™ assays to confirm the binding of all compounds in MLLT1. In agreement, the measured IC50 values for all compounds correlated well with their KD values obtained from ITC (Figure 4A and B). In essence, **1** was the most potent binder among the piperazine-urea derivatives, and its low micromolar potency was similar to that observed for the benzimidazole-amide counterparts. In addition, we observed weak binding of these compounds to MLLT3, and much lower affinities or no detectable activity for YEATS2 and YEATS4 (Supplementary table S2). In general, a trend of lower affinities of piperazine-urea compounds than the benzimidazole-amide derivatives could be due to the lack of potential aromatic stacking with Phe28 and Phe59. Nevertheless, the different binding mode of the piperazine-urea compounds indicated a possibility of targeting the previously unseen rear pocket towards loop 4. Hybrid compounds between these two scaffolds exploiting larger area of the rear pocket as well as maintaining aromatic stacking with Phe28 and Phe59 might offer a strategy for potent inhibitors for targeting MLLT1.

**Figure 4.**
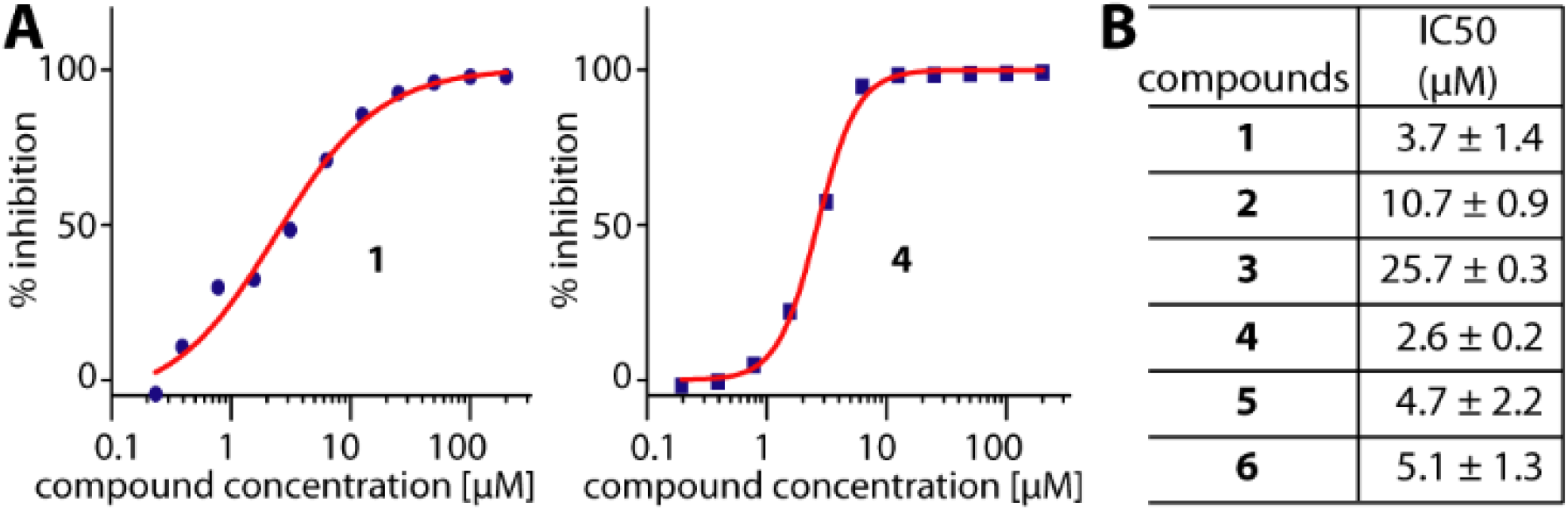
AlphaScreen™ assays for inhibition of MLLT1. A) Examples of MLLT1 inhibition with **1** and **4**. B) Summary of IC50s.

YEATS-domain-containing proteins plays a key role in development of many diseases, presenting a potential target for chemotherapeutic treatment. Although sharing similar functions in recognition of acetyl-lysine, targeting this reader family requires distinct classes of small molecule binders to those for bromodomains. Our previous effort has identified benzimidazole-amide as an acetyl-lysine mimetic scaffold for MLLT1/3, which led to development of the first chemical probe for MLLT1/3, albeit with limited pharmacokinetic properties. Here we characterized another chemotype in piperazine-urea, of which the derivatives demonstrated also affinities in low micromolar range for MLLT1. Comparative structural analyses suggested that this alternative scaffold likely exhibited different interaction mechanisms, and exploited diverse binding cavities. The piperazine-urea scaffold not only presents an alternative chemical starting point, but expands the existing knowledge on landscape of potential druggable pockets of the protein, which would benefit future programs on the development of inhibitors for this attractive epigenetic targets.

## EXPERIMENTAL SECTION

Piperazine-urea compound **1-3** were purchased from Enamine^25^, while benzimidazole-amide derivative **4-6** were obtained from our previous study^26^.

Protein purification and crystallization: MLLT1 YEATS domain (aa 1-148) was purified using the procedure described previously^21^. Apo crystals were produced using sitting drop vapour diffusion method at 20 °C and the condition containing 25% PEG 3350, 0.2 M NaCl, and either 0.1 M bis-tris, pH 6.5 or 0.1 M HEPES, pH 7.5. Ligands were soaked into crystals. Diffraction data were collected at SLS X06SA, processed with XDS^31^ and scaled with aimless^32^. Molecular replacement was performed with Phaser^33^ and the MLLT1 apo structure (pdb-id 6hq0). All structures were rebuilt in COOT^34^, refined using REFMAC^35^, and validated with molprobity^36^. Data collection and refinement statistics are summarized in Supplementary table S1.

Isothermal calorimetry (ITC) experiment were performed using NanoITC instrument (TA instruments) at 25°C with protein at 300 μM in 25 mM Tris pH 7.5, 500 mM NaCl, 0.5 TCEP, 5% glycerol titrating into 20-30 μM compounds. Data analyses were carried out using the software and protocols provided by TA instruments.

AlphaScreen™ assays for all compounds with MLLT1 were performed as previously described^25^.

## AUTHOR INFORMATION

### Author Contributions

XN, DH, TC and JM conducted the experiment. MM and PB contributed inhibitors. AC, SK, OF, PB supervised research. AC and SK wrote the paper. All authors have given approval to the final version of the manuscript.

### Notes

Coordinates and structure factors have been deposited with accession codes 6t1i, 6t1j, 6t1l, 6t1m, 6t1n and 6t1o.

## ACKNOWLEDGMENT

The SGC, a registered charity that receives funds from AbbVie, Bayer Pharma AG, Boehringer Ingelheim, Canada Foundation for Innovation, Eshelman Institute for Innovation, Genome Canada, Innovative Medicines Initiative [ULTRA-DD 115766], Wellcome Trust, Janssen, Merck & Co., Novartis Pharma AG, Ontario Ministry of Economic Development and Innovation, Pfizer, São Paulo Research Foundation-FAPESP, Takeda and the Centre of Excellence Macromolecular complexes (CEF). S.K. and A.C. are supported by the Sonderforschungsbereich SFB1177 Autophagy. M.M. is grateful to the EPSRC Centre for Doctoral Training in Synthesis for Biology and Medicine (EP/L015838/1) for a studentship, supported by AstraZeneca, Diamond, Defence Science and Technology Laboratory, Evotec, GlaxoSmithKline, Janssen, Novartis, Pfizer, Syngenta, Takeda, UCB and Vertex. The authors thank staffs at Swiss Light Source for their assistance during data collection, which has been supported also by the funding from the European Union’s Horizon 2020 research and innovation program under grant agreement number 730872, project CALIPSOplus.

## ABBREVIATIONS

YEATS domain: The Yaf9, ENL, AF9, Taf14, Sas5 (YEATS) domain
ENL: eleven-nineteen-leukemia protein
MLLT1: mye-loid/lymphoid or mixed-lineage leukemia translocated to, chromosome 1 protein
AF9: ALL1-fused gene from chromosome 9 protein
MLLT3: myeloid/lymphoid or mixed-lineage leukemia translocated to chromosome 3 protein

## Supplementary Information

**Supplementary table S1.**
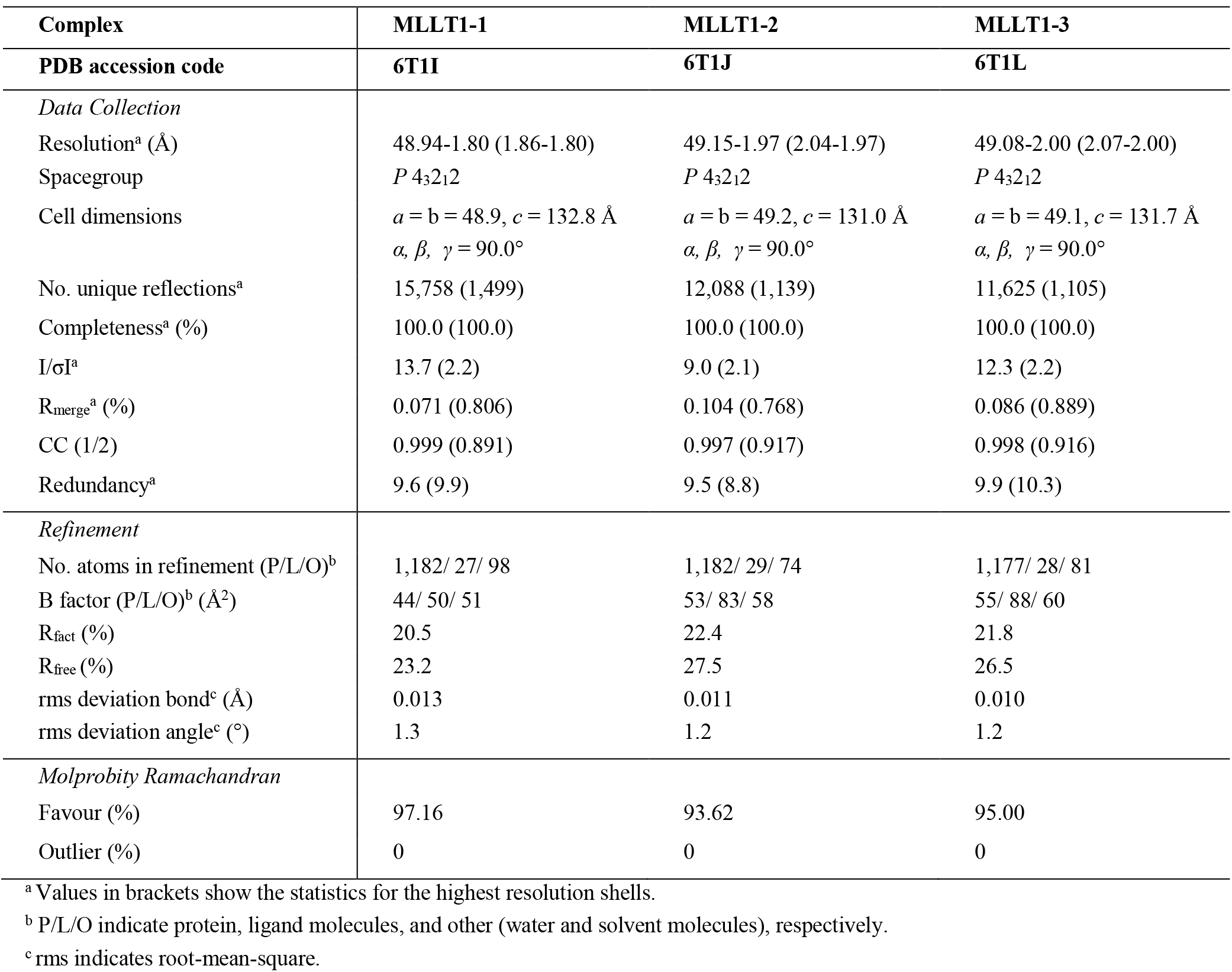
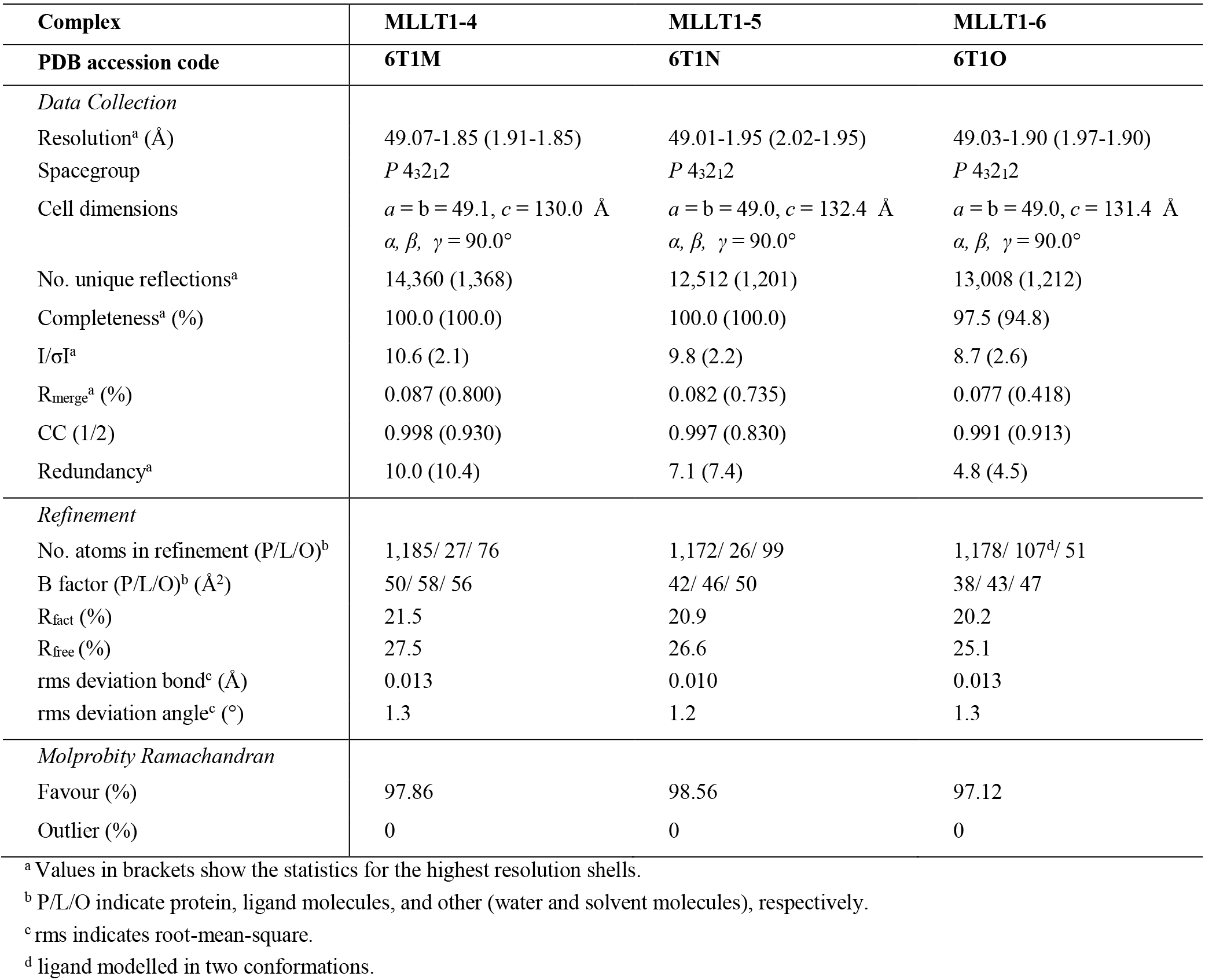
Data collection and refinement statistics

**Supplementary table S2.** Summary of IC50s of the compounds against other YEATS domains.

| compound | IC50 ($\mu\text{M}$) | | |
| --- | --- | --- | --- |
|  | MLLT3 | YEATS2 | YEATS4 |
| 1 | 7.8 | 76.8 | >100 |
| 2 | 33.0 | 32.2 | >100 |
| 3 | >100 | >100 | >100 |
| 4 | n.d | >100 | >100 |
| 5 | n.d | 53.2 | >100 |
| 6 | n.d | 60.4 | >100 |

**Supplementary figure S1.**
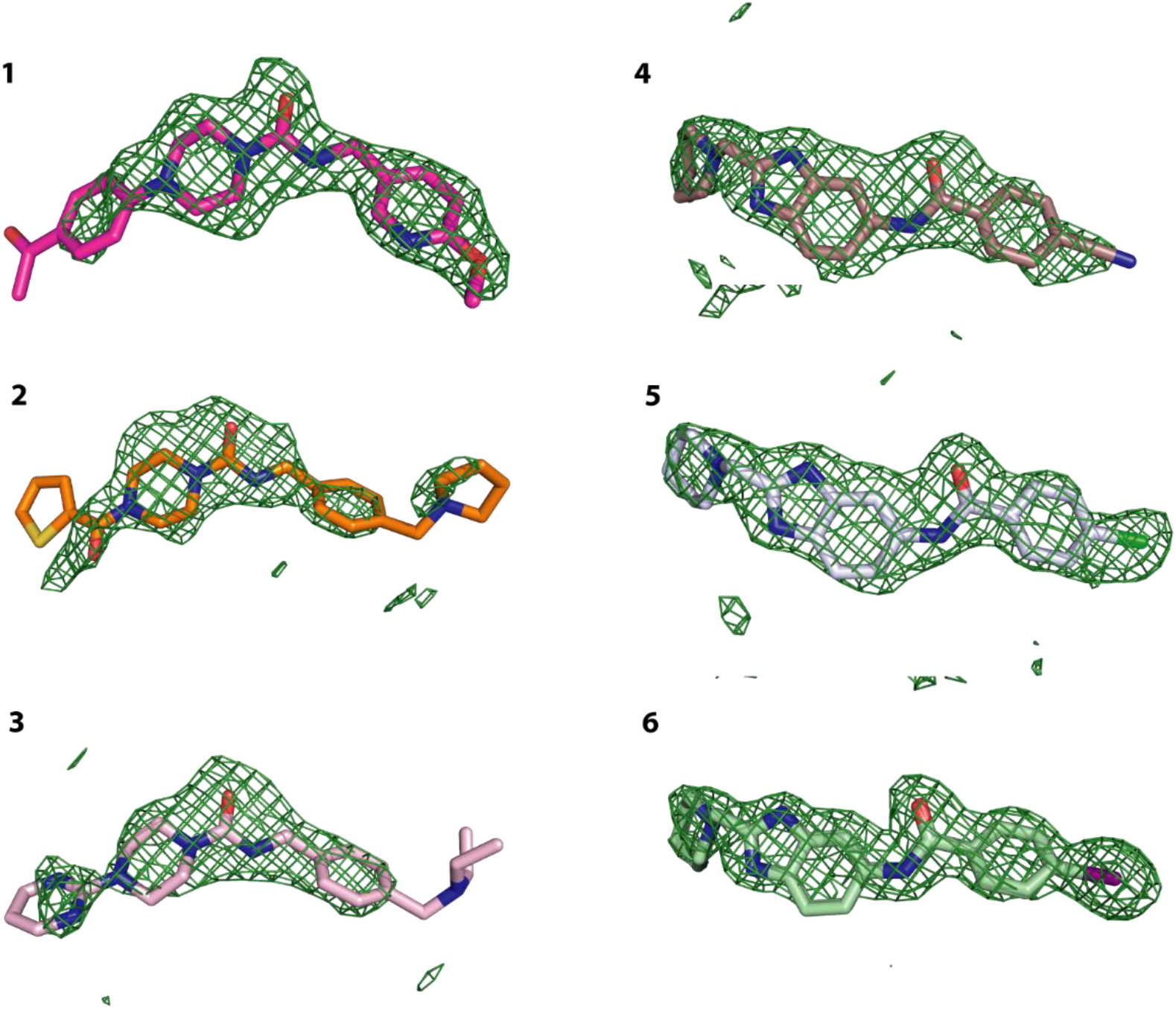
Omitted electron density maps for the bound ligands. The |F_O_|-|F_C_| electron density maps contoured at 2.5σ.

